# Aging reveals divergent responses of AgRP/NPY neurons to diet in male and female mice

**DOI:** 10.1101/2025.11.20.689576

**Authors:** Tionna Brothers, Wei Wei, Austin C. Korgan, Zoey Bridges, Kristen M.S. O’Connell

**Affiliations:** The Jackson Laboratory for Mammalian Genetics, Bar Harbor ME USA; Tufts University Graduate School of Biomedical Sciences, Boston MA USA

**Keywords:** hypothalamus, diet-induced obesity, sex differences, energy balance, inflammation, patch-clamp

## Abstract

**Objectives:** Neurons coexpressing Agouti-related peptide (AgRP) and Neuropeptide Y (NPY) are an essential component of an interoceptive circuit regulating hunger and metabolism. Their activity is closely linked to metabolic state and their output is sensitive to diet-induced plasticity, which may influence the development of obesity and associated metabolic diseases. However, most studies use young male mice, even though obesity and its comorbidities are sensitive to both biological sex and aging, leaving a significant gap in our understanding of the role of these neurons in females and in older animals. Our goal was to begin to address this gap by investigating the effects of diet and age on AgRP/NPY neuronal activity in female mice in both early adulthood and midlife.

**Methods:** Female transgenic NPY-GFP mice aged 8 – 32 weeks were fed either a standard control chow diet or a high-fat, high-sugar diet (HFD) for 8 – 24 weeks and brain slice patch clamp electrophysiology was used to measure the response of AgRP/NPY neurons.

**Results:** We found that in young, lean female mice, the baseline firing rate of AgRP/NPY neurons is significantly elevated compared to age-matched males, thus the impact of HFD on the output of these neurons is blunted relative to control. However, in the baseline firing rate of neurons from lean middle-aged female mice is significantly lower, resulting in a greater relative impact of HFD on AgRP/NPY neuronal output, the development of neuronal leptin resistance, and significant weight gain.

**Conclusions:** Both sex and age significantly impact the function and modulation of AgRP/NPY neurons, emphasizing the need to include these biological variables in experimental design.

**Highlights:**

- Male and female mice exhibit sex-dependent responses to a high-fat diet, with female mice requiring twice as long to develop an overt obese phenotype.
- AgRP/NPY neurons in young, lean female mice have significantly higher activity than those from male mice.
- Aging female mice develop obesity and altered function and leptin sensitivity of AgRP/NPY neurons.
- The hypothalamic glial response to high-fat diet is influenced by sex – age interactions.

## 1. INTRODUCTION

Located in the arcuate nucleus of the hypothalamus (ARH), neurons coexpressing agouti-related peptide (AgRP) and neuropeptide Y (NPY) are key regulators of metabolism and energy expenditure [1–3]. These neurons play a key role in the integration of central and peripheral signals to regulate food intake and body weight [4–7]. AgRP/NPY neurons also influence a wide range of other behaviors, including circadian regulation of feeding [8], motivation to exercise [9; 10], cortical development [11], hepatic glucose regulation [12], nutrient partitioning [13], and thermogenesis [14]. Prior work has described alterations of body weight and metabolism as dependent on AgRP neuronal function and expression [15–19], particularly AgRP neuronal firing rate in response to fasting [20–22], acute [7] or long-term HFD [23–26], and sugar ingestion [27; 28].

Studies on AgRP neuron activity related to food intake and energy metabolism are based predominantly on data from young (<6 months old) male mice [1]. Sexual dimorphism in metabolism and energy expenditure is well characterized [29–35] but there is a gap in our understanding of the effects of HFD on hypothalamic circuits in the context of aging and female sex. Previous reports suggest that female mice are resistant to diet-induced weight gain using experimental paradigms that robustly induce obesity in in male mice [3]; an approach optimized for inducing DIO in female mice has yet to be developed. Similarly, there has been little attention paid to the impact of obesity in aging individuals, despite the fact that mid-life obesity increases risk for a number of age-related diseases, including type 2 diabetes, cardiovascular disease, Alzheimer’s disease, and others [35–38]. Thus, a better understanding of the impact – and interaction - of sex and age in CNS mediated control of food intake behaviors and maintenance of weight gain during development of diet-induced obesity is essential to develop better preclinical models for translational studies that inform future healthcare interventions.

Here, we show that AgRP neurons from young, lean female C57BL/6J mice have a higher baseline firing, which mitigates the effect of HFD feeding on the intrinsic excitability of AgRP/NPY neurons compared to age-matched males; these young female mice also fail to develop DIO or neuronal leptin resistance. We further show that age plays a role in the activity of AgRP/NPY neurons, as baseline firing of these neurons from middle-aged female mice is significantly decreased, resulting in a more pronounced effect of HFD on neuronal excitability, which is associated with the development of obesity and a failure of leptin to inhibit the activity of AgRP/NPY neurons. These findings represent a divergence between hypothalamic activity and metabolism in female mice that has not been previously characterized.

## 2. MATERIALS AND METHODS

### 2.1. Animals

All animal care and experimental procedures were approved by The Animal Care and Use Committee at The Jackson Laboratory. Electrophysiology experiments used both male and female mice from the transgenic strain B6.FVB-Tg(NPY-hrGFP)1Lowl/J (NPY-GFP; JAX Strain#: 006417). Immunohistochemistry experiments used male and female C57BL/6J (B6; JAX Strain#: 000664). Mice were housed at 22-24° C on a 12h:12h light/dark cycle (0600 - 1800) and were fed standard lab chow (LabDiets 5K0Q: 3.15 kcal/g metabolizable energy, 16.8 kcal% from fat). At 8 weeks of age, experimental mice were randomly assigned to either normal chow diet (NCD) or to high-fat diet (HFD; Research Diets D12451: 4.73 kcal/g metabolizable energy, 45 kcal% from fat) for 8 weeks (young cohort) or 24 weeks (middle-aged cohort; MA). Water and food were available *ad libitum*. Mice were weighed weekly and used for experiments at ∼16 and 32 weeks of age.

### 2.2. Electrophysiology

#### 2.2.1. Slice preparation

Slice electrophysiology was performed as previously described [24; 39]. For all experiments, brain slices were prepared between 0900 and 1030. Mice used for electrophysiology experiments were deeply anesthetized using isoflurane prior to decapitation and rapid removal of the brain. The brain was immediately submerged in ice-cold, oxygenated (95% O_2_/5% CO_2_) cutting solution (in mM: 119 NaCl, 90 Sucrose, 2.5 KCl, 1 MgSO_4_, 2 CaCl_2_, 1.25 NaH_2_PO_4_, 23 NaHCO_3_, and 10 glucose). Coronal slices (250μm) were cut using a vibratome (VT1000S, Leica) and incubated in oxygenated aCSF (in mM: 119 NaCl, 2.5 KCl, 1 MgSO_4_, 2 CaCl_2_, 1.25 NaH_2_PO_4_, 23 NaHCO_3_, and 10 glucose) for 1h prior to recording.

#### 2.2.2. Slice recording

Slices were transferred to a recording chamber constantly perfused (∼2 ml/min) with oxygenated aCSF. GFP-positive NPY neurons were identified using epifluorescence and standard GFP filters on a fixed-stage Scientifica (Uckfield, UK) SliceScope 1000 microscope equipped with a digital camera (Q-Imaging, Surry, BC, Canada). All recordings were performed using a Multiclamp 700B amplifier and Digidata 1550A, controlled using Clampex 10.7 (Molecular Devices, San Jose, CA, USA). Data were digitized at 20 kHz and filtered at 5 kHz using the built-in four-pole Bessel filter of the Multiclamp 700B.

Recording pipettes were pulled from filamented thin-wall borosilicate glass (TW150F-4, World Precision Instruments) and had a resistance of 4-7 MΩ when filled with internal solution (for intrinsic excitability (AP) recordings, in mM: 130 K-gluc, 10 KCl, 0.3 CaCl_2_, 1 MgCl_2_, 1 EGTA, 3 MgATP, 0.3 NaGTP, and 10 HEPES, pH 7.35 with KOH). The liquid junction potential (LJP) between normal aCSF and the K-gluconate solution used for intrinsic recordings was +14.7 mV and was corrected.

Whole-cell current clamp recordings of the resting membrane potential (RMP) and spontaneous action potential firing were performed in the presence of DNQX (10 μM, Tocris) and picrotoxin (100 μM, Tocris). For experiments testing inhibition of AgRP/NPY neurons by leptin, NCD mice were fasted overnight to promote increased intrinsic excitability (HFD mice were not fasted); following ∼2 minutes of baseline recording, 100 nM leptin (Tocris) was bath applied.

#### 2.2.3. Data analysis and statistics

Current clamp recordings were analyzed in Clampfit 10.7 (Molecular Devices). Group differences were analyzed with two- or three-way ANOVA followed by Tukey’s multiple comparisons *post hoc* comparisons using the *aov* and *TukeyHSD* functions in R (4.5.0). For repeated measures analysis, group differences were analyzed by two-way RM-ANOVA followed by pairwise comparisons using Bonferroni corrected t-tests. Data visualization was performed using r/ggplot in R or GraphPad Prism. For all statistical tests, a value of *p* < 0.05 was considered significant. Data are presented as the mean ± SEM.

### 2.3. Immunohistochemistry

Mice were transcardially perfused with phosphate-buffered saline (PBS) for ∼5 min, followed by 4% paraformaldehyde in 1x PBS solution for ∼10 min. Brains were extracted and postfixed overnight in 4% Paraformaldehyde in 1x PBS at 4°C. The following day, brains were transferred to 1x PBS until they were coronally sectioned at 75-100 µm using a vibratome (VT1000S, Leica).

#### 2.3.1. Primary Antibodies

Brains were blocked for two hours at room temperature in blocking solution (0.1% TritonX + 5% normal donkey serum + 3% BSA in PBS), then transferred to blocking solution + primary antibody for two nights at 4°C. Primary antibodies used were chicken anti-GFAP (Abcam Ab4674, 1:1000) and goat anti-Iba1 (Abcam Ab5076, 1:1000).

#### 2.3.2. Secondary antibodies

Brains were washed three times with PBS before incubation overnight at 4°C in secondary blocking solution that included secondary antibodies donkey anti-chicken AlexaFluor488 (1:2000; Thermo A11055), donkey anti-goat AlexaFluor647 (1:2000; Jackson Immunology 703-605-155), and 1:2000 DAPI (ThermoScientific 62248). Sections were washed three times with PBS, mounted on slides, and cover-slipped with Fluoromount-G (ThermoScientific 00-4958-02).

#### 2.3.3. Image Analysis

Images were taken using a STELLARIS 5 (Leica Microsystems) laser scanning confocal microscope, acquired with Leica Application Suite X (LAS-X, 3.7.6.25997), and processed with Fiji/ImageJ software. Fluorescence intensity was quantified by measuring the raw integrated density in FIJI and normalizing to the total area measured per image stack.

### 2.4. Data Availability

All relevant data are available upon request from the authors.

## 3. RESULTS

### 3.1. Sex and age influence the response to HFD in female mice

To determine the impact of HFD on body weight, AgRP/NPY neuronal excitability, and leptin sensitivity in female mice using a common experimental paradigm for DIO in male mice, we randomly assigned 8-week old female mice to either a normal chow diet (NCD) or a high-fat/high-sugar (HFD) diet for 8 weeks. As expected, female mice are significantly smaller than their age-matched male counterparts, with sex accounting for the greatest proportion of variation in body weight (45.7%) across the study population (*F_(1,75)_ =* 179.1, p < 0.0001). Consistent with prior reports, male mice fed HFD develop significant weight gain within 2 weeks of switching diets (*F_(1,37)_* = 14.38, p < 0.0005, Figure 1A). In contrast, age-matched female mice fed HFD exhibit almost no weight gain over the same time frame, with no significant main effect of diet (*F_(1, 38)_ =* 2.724). There were no significant differences between the NCD- and HFD-fed female cohorts at any time point in this 8 week study, consistent with previous reports (Figure 1A) [32; 40; 41].

**Figure 1:**
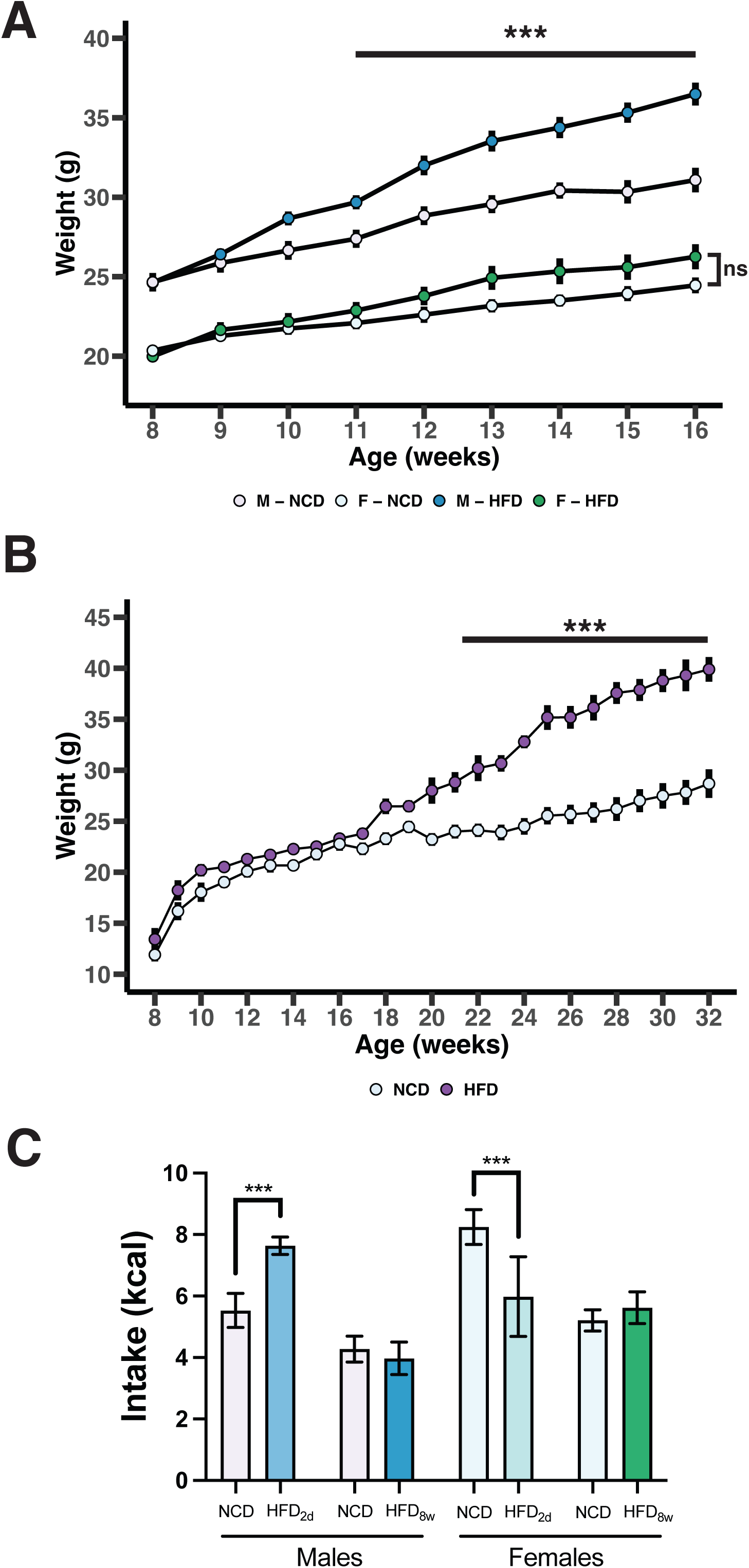
Age and sex interact to influence the onset of diet-induced obesity in mice. **(A)** Body weight curve for male and female mice fed either NCD or HFD for 8 weeks. (**B)** Body weight curve for female mice fed either NCD or HFD for 24 weeks beginning at 8 weeks of age. (**C)** Food intake measured over a 24h period 2d and 8 weeks after beginning HFD. Data expressed as mean ± SEM. For Panels A and B, ***p < 0.001, repeated measures ANOVA. For Panel C, ***p < 0.001, unpaired t-test compared to group-matched NCD.

In humans, the prevalence of obesity is roughly equal in men and women, although obesity and its comorbidities often present differently in each sex [42; 43]. Thus, while female mice are often described as ‘resistant’ to DIO, we explored the possibility that protocols for DIO require optimization for application to female populations and that, given the sexual dimorphism in metabolic pathways, female mice may require a longer schedule of HFD feeding to facilitate the development of an obese phenotype. We therefore started a separate cohort of female mice on either NCD or HFD and continued feeding the assigned diet until we observed a significant change in body weight between the two groups. As shown in Figure 1B, as in the young cohort, female mice on HFD did not exhibit body weight gain over the first 14 weeks of HFD feeding, but starting around week 15, they begin to diverge from their age-matched NCD-fed counterparts and continue to diverge from their age-matched NCD-fed counterparts for the remainder of the study. These data demonstrate that female mice are indeed susceptible to diet-induced obesity but require nearly twice as long to develop the phenotype as age-matched male mice.

Previous reports have shown that male mice will transiently increase their caloric intake upon consuming a HFD, though intake eventually normalizes to be roughly equivalent to age- and sex-matched mice fed a low-fat control diet [44]. To determine if female mice display a similar diet-induced hyperphagia, we measured 24h food intake 2 days or 8 weeks after switching male and female mice to HFD. As shown in Figure 1C, as expected, male mice initially increase their energy intake significantly compared to NCD-fed controls, and after 8 weeks of HFD feeding reduce intake to levels comparable to age-matched mice that have never eaten HFD. Female mice, on the other hand, initially exhibit the opposite response to HFD and significantly reduce caloric intake compared to age- and sex-matched controls when measured 2 days after introducing HFD; similar to male mice, female mice also normalize calorie intake to a level comparable to controls after 8 weeks of HFD feeding.

### 3.2. Sex dimorphism in the activity of AgRP/NPY neurons in young female mice

Given the well-known role of AgRP/NPY neurons in both body weight and regulation of food intake, we next determined whether the activity and modulation of these neurons is influenced by sex. We previously demonstrated that in male mice fed HFD, even for very brief periods, AgRP/NPY neurons become hyperexcitable and refractory to modulation by physiological cues of hunger and satiety, even before the onset of significant weight gain [3; 24]. We therefore hypothesized that the apparent resistance of female mice to DIO following short-term (8 weeks) HFD feeding may be accompanied by a similar resistance to diet-induced remodeling of AgRP/NPY neuronal excitability. We used whole-cell patch clamp to measure the intrinsic properties of AgRP/NPY neurons from female mice fed either NCD or HFD for 8 weeks. There was a significant main effect of sex on the baseline activity of AgRP neurons in lean NCD_fed_ mice – neurons from satiated male mice exhibited the expected low rate of spontaneous activity, but neurons from NCD_fed_ female mice fired at a significantly faster rate compared to those from male mice (NCD_fed_ males: 1.1 ± 0.17 s^-1^, NCD_fed_ females: 2.6 ± 0.52 s^-1^, p = 0.008, Figure 2A). As expected, overnight fasting significantly increased the output of AgRP/NPY neurons from male mice (NCD_fast_ males: 3.4 ± 0.38 s^-1^, p < 0.0001); overnight fasting increased the firing of AgRP/NPY neurons from female mice to a comparable rate to male mice (NCD_fast_ females = 3.8 ± 0.56 s^-1^, p = 0.47) but this was not significantly higher than the baseline rate measured in NCD_fed_ female mice (NCD_fed_ vs NCD_fast_ females: p = 0.08). Similar to what we previously reported in male mice [3; 24] (Figure 2B), HFD feeding increased the spontaneous firing rate of AgRP/NPY neurons from female mice (HFD females: 4.5 ± 0.45 s^-1^, p = 0.002 vs. NCD_fed_); notably, the diet-associated change in AgRP/NPY neuronal output in female mice is significantly greater than that observed in male mice (HFD males: 3.04 ± 0.33, p = 0.002 vs female HFD). As an additional measure of intrinsic excitability, we also measured the resting membrane potential (RMP), which we have previously reported to be depolarized by both fasting and HFD in male animals [3; 24] and found that in females, HFD – but not fasting – was associated with a slight, but significant depolarization of the RMP (Figure 2C; NCD_fed_ = - 60.0 ± 0.6 mV; NCD_fast_ = -58.9 ± 0.8 mV; HFD = -56.3 ± 0.6 mV, *F_(2,52)_* = 8.991, p = 0.0004).

**Figure 2:**
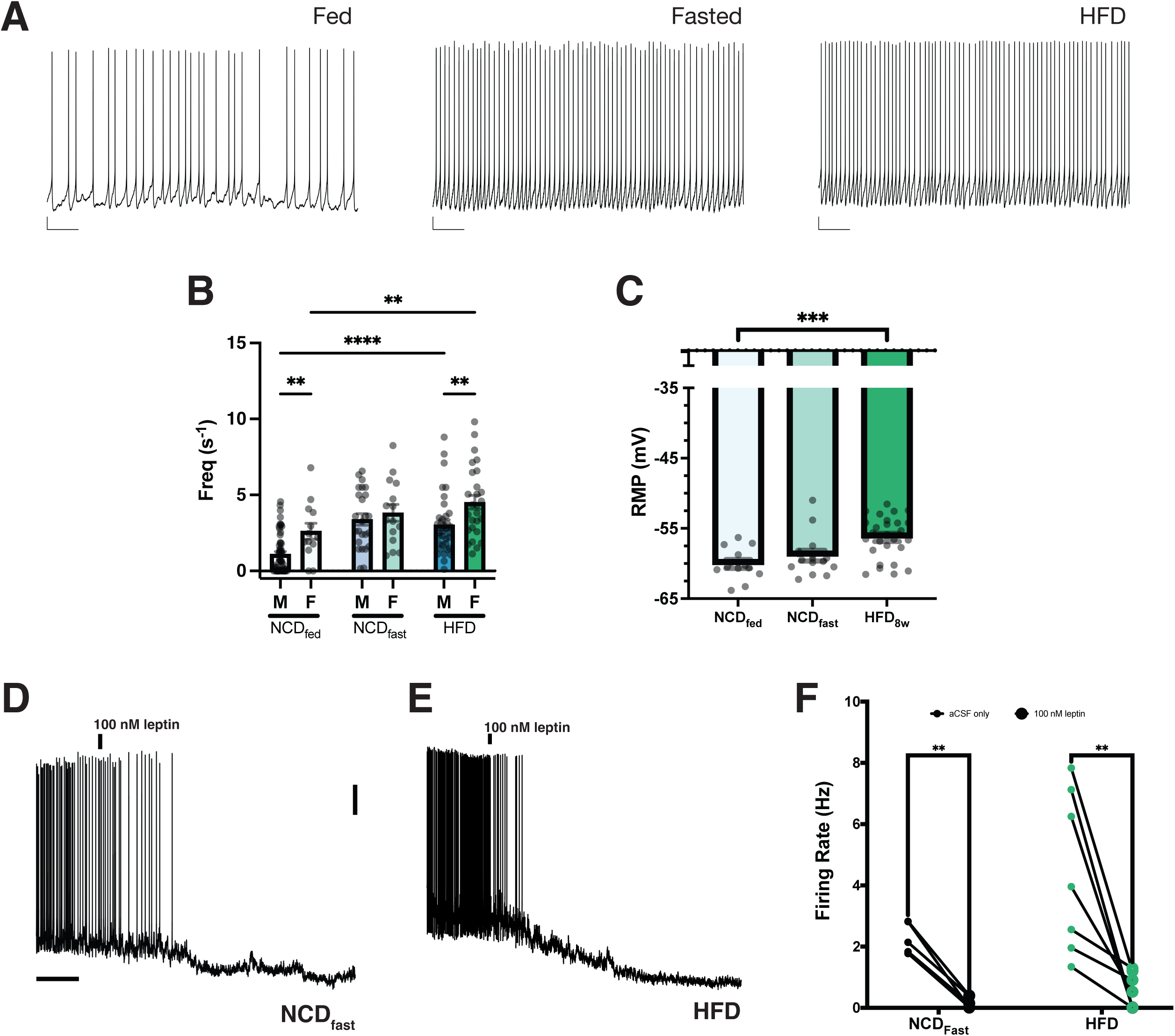
AgRP/NPY neuronal function and response to high-fat diet is sexually dimorphic in young mice. **(A)** Representative current-clamp recordings of spontaneous action potentials from AgRP/NPY neurons. Scale: vertical = 5 mV, horizontal = 10 sec. (**B)** Mean spontaneous AP frequency from current-clamp recordings of AgRP/NPY neurons at baseline (NCDfed), following 16h fast (NCDfast), and 8 weeks of HFD (HFD) in age-matched male and female mice. (**C)** Resting membrane potential of neurons in (**B**) Representative current-clamp recordings of AgRP/NPY neurons from either NCDfast (**D**) or HFD (**E**) female mice during bath application of 100 nM leptin. (**F)** Summary plot of the AP frequency before and after addition of leptin. For Panels B and C, **p < 0.01, ***p < 0.001, ****p <> 0.0001, Bonferroni-adjusted p-values. For panel F, **p < 0.001, paired t-test.

Given that male mice fed HFD for 8 weeks develop hypothalamic leptin resistance [3; 45; 46] and the presence of sex differences in the function and plasticity of AgRP/NPY neurons, we next assessed whether the response of these neurons to leptin is affected by 8 weeks of HFD feeding. As in male mice, bath application of 100 nM leptin significantly inhibits the firing of AgRP/NPY neurons in brain slices from lean, fasted NCD female mice (aCSF only: 2.3 ± 0.2 s^-1^; + leptin: 0.2 ± 0.1 s^-1^, paired *t_5_* = 9.204, p = 0.002) (Figure 2D and E). In contrast to our previous findings in male mice, AgRP/NPY neurons from female mice fed HFD for 8 weeks remained highly sensitive to leptin (HFD aCSF: 4.4 ± 0.6 s^-1^; +leptin: 0.6 ± 0.2 s^-1^, paired *t_6_* = 3.73, p = 0.01; Figure 2F).

### 3.3. Effect of age on the neuronal response to HFD in female mice

In humans, the rates of overweight and obesity in males and females are comparable, though there are known sex differences in the development of obesity and associated comorbidities[47]. Thus, the common perception that female mice are resistant to diet-induced obesity is likely a consequence of the under-utilization of female subjects in biomedical research. Since we observed significant sex differences in the function and plasticity of AgRP/NPY neurons, we reasoned that DIO paradigms that efficiently result in DIO in males may not be optimized for females. Therefore, we extended our DIO paradigm to determine the duration of HFD feeding required to establish a model of obesity in female mice. As before, a cohort of female NPY-GFP mice were randomly assigned to either NCD or HFD at 8 weeks of age, but were continued on the assigned diet until HFD mice began to significantly weigh more than their age-matched NCD controls. As shown in Figure 1B, on the same 45kcal% diet used in young mice, female mice require twice as long – 16 weeks – of HFD feeding before developing an obese phenotype, clearly demonstrating that female mice are not intrinsically resistant to DIO, rather they exhibit a delayed response to diet. For consistency with experiments using male mice, once female mice began to gain weight, they were maintained on HFD for another 8 weeks before being used for electrophysiological assessment of AgRP/NPY neuronal function.

To determine whether the excitability of AgRP/NPY neurons is altered in middle-aged (MA, 24 weeks of age) female mice, we again used brain slice electrophysiology to assess baseline function of these neurons in lean, MA NCD female mice. As shown in Figure 3A and B, the baseline firing rate of AgRP neurons from 8 month old MA female mice is significantly lower than what we observed in young (4 month old) female mice (MA: 1.2 ± 0.32 s^-1^, adj. p = 0.01 vs. young females; see Figure 2) and is more similar to that measured in young male mice (adj. p = 0.56). This age-dependent change in AgRP/NPY neuronal firing is due principally to a change in the distribution of firing rates across the entire population of recorded neurons, with an increase in ‘silent’ neurons in MA females relative to young females, while the maximal observed firing rate was unchanged. In both young male and MA female mice, >50% of AgRP/NPY neurons from lean, NCD-fed animals fire at a rate of < 1 s^-1^ (p = 0.57, Fisher’s exact test), while in young NCD-fed females, significantly fewer - only ∼15% - AgRP/NPY neurons are silent at baseline (p = 0.0008 vs young males, p = 0.01 vs MA females, Fisher’s exact test).

**Figure 3:**
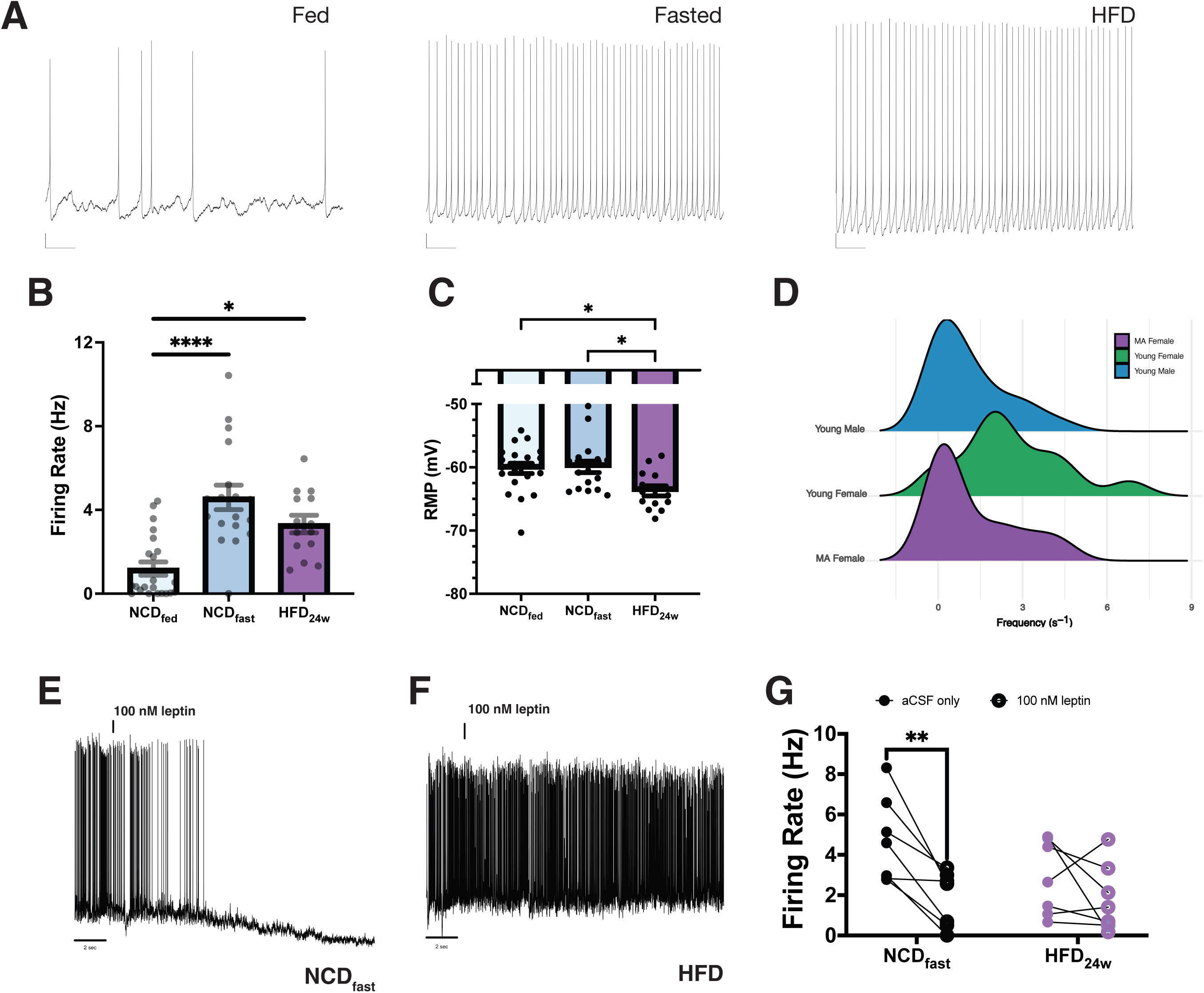
AgRP/NPY neuronal function and leptin sensitivity is altered in middle-aged female mice. **(A)** Representative current-clamp recordings of spontaneous action potentials in AgRP/NPY neurons from 8 month old female mice. Mean spontaneous AP frequency (**B**) and RMP (**C**) of AgRP/NPY neurons from baseline (NCDfed), 16h fasted (NCDfast), and HFD (HFD) female 8 month old mice. (**D**) Ridgeline plots illustrating the impact of age and sex on the distribution of AP frequencies in AgRP/NPY neurons. Representative current-clamp recordings of AgRP/NPY neurons from middle-aged NCDfast (**E**) and HFD (**F**) female mice before and after bath application of 100 nM leptin. (**G**) Summary plot of the AP frequency before and after addition of leptin. For panels B and C, *p < 0.05, ****p < 0.0001, Bonferroni-adjusted p-values. For panel G, **p < 0.001, paired t-test.

We next determined whether AgRP/NPY neurons from MA female mice fed HFD for 24 weeks developed leptin sensitivity. There was no significant main effect of age alone on leptin inhibition of AgRP/NPY neurons, as bath application of 100 nM leptin resulted in significant inhibition of neuronal activity (Figure 3D and E). However, in contrast to what we observed in young female HFD mice, AgRP/NPY neurons from MA female mice on HFD for long enough to develop DIO were no longer inhibited by leptin, indicating that the diet-induced hyperexcitability of AgRP/NPY neurons in MA DIO mice is accompanied by the development of hypothalamic leptin resistance (Figure 3F and G).

### 3.4 Glial Response to High Fat Diet in the Hypothalamus

Hypothalamic inflammation and gliosis are commonly observed in both humans with obesity [48–50] and rodent models of obesity [51–56], suggesting a potential causal relationship between hypothalamic gliosis and the development of obesity. Although most rodent studies predominantly use male animals, a few reports using both sexes suggest that there are sex differences in the development of hypothalamic inflammation and both astrogliosis and microgliosis [57; 58]. To determine if there are sex differences in the response of astrocytes and microglia in our model of DIO, we used immunofluorescence staining against either GFAP or Iba1 in the mediobasal hypothalamus (MBH), which includes the ARH, of male and female mice fed HFD for either 8 or 24 weeks to match the HFD feeding schedule used in our electrophysiology experiments. As shown in Figure 4A - C, there was no significant difference in the fluorescence intensity of MBH GFAP with either age or diet in female mice. On the other hand, male mice (Figure 4A-C) have increased MBH GFAP immunoreactivity with both age and diet; the levels of GFAP-IR in male HFD mice are also significantly higher than in age- and diet-matched female mice, suggesting that there is sex dimorphism in astrocyte responsiveness in the MBH. To evaluate the effect of age, sex, and diet on microglial activation, we used immunofluorescence staining against Iba1, a commonly used marker for activated microglia. In both male and female mice, there was a significant increase in Iba1-IR with age (*F_1,31_* = 128.54, p < 0.000001). Female mice exhibited no significant effect of diet on Iba1-IR at either age (Figure 4D-F). Young (4 months old) male DIO mice had a non-significant trend toward higher Iba1-IR in the MBH (adj. p = 0.08) that was significant in middle-aged male mice (adj. p = 0.02), suggesting that there are sex- and age-interactions in both astrocytic and microglial response to HFD.

**Figure 4:**
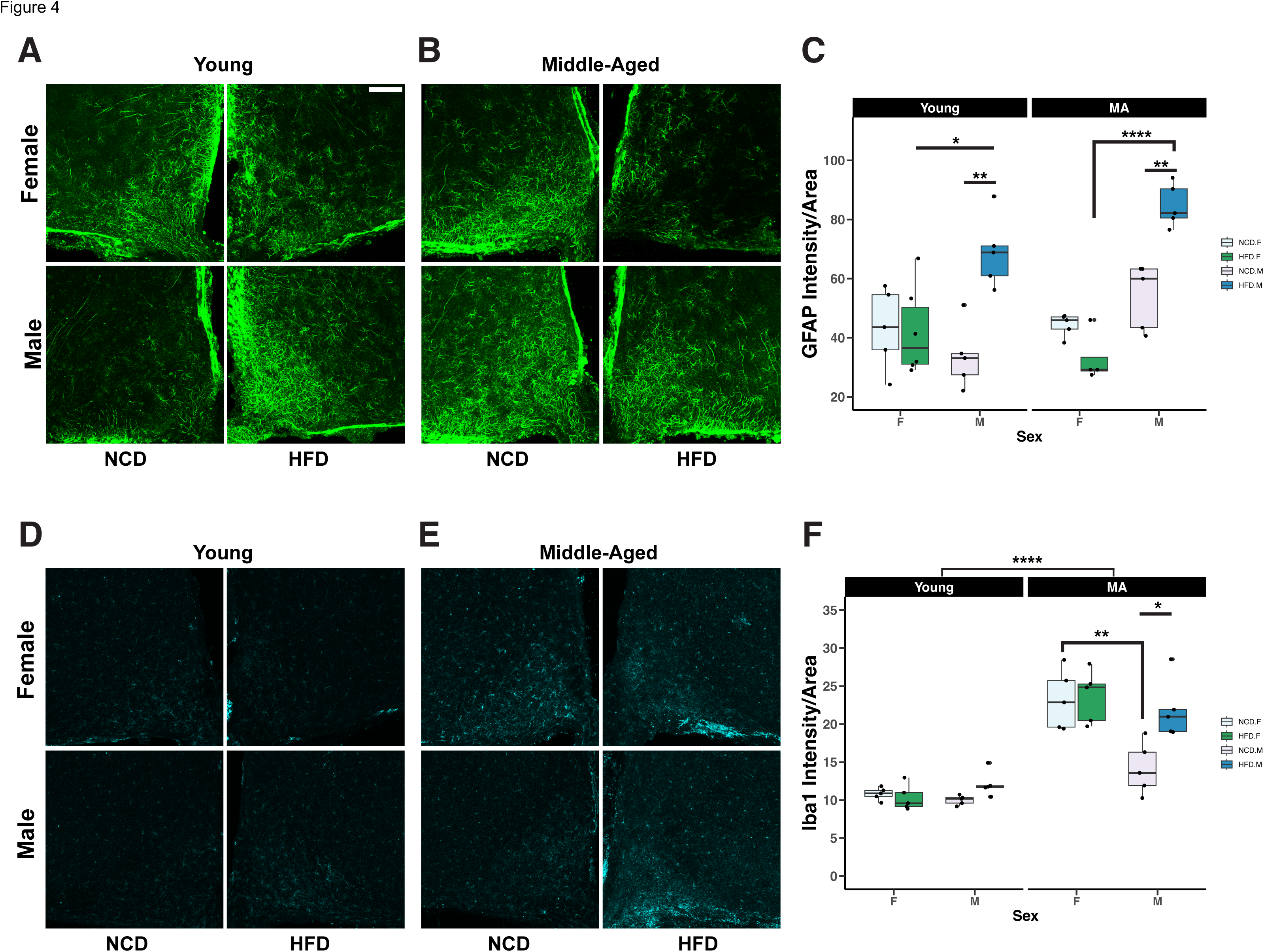
Sex, age, and diet influence the morphological response of astrocytes and microglia in the mediobasal hypothalamus. (**A**) Representative images of GFAP-positive astrocytes in the MBH from young male and female mice. (**B**) Representative images of GFAP-positive astrocytes in the MBH of middle-aged male and female mice. (**C**) Summary boxplot of the GFAP integrated fluorescence density normalized to total area. (**D**) Representative images of Iba-1 positive microglia in the MBH of young male and female mice. (**E**) Representative images of Iba-1 positive microglia from the MBH of middle-aged male and female mice. (**G**) Summary boxplot of the Iba1 integrated fluorescence density normalized to total area. Scale bar = 100 μm and applies to all images. *p < 0.05, **p < 0.01, ****p < 0.0001, three-way (Diet * Sex * Age) ANOVA with TukeyHSD *post hoc* corrections.

## 4. DISCUSSION

Continued focus on the maintenance of body weight as it relates to health outcomes like type 2 diabetes [59–62], combined with increased popularity of pharmacological interventions for weight [63] necessitate further study of how the central nervous system regulates and cooperates with peripheral systems that maintain food intake and energy expenditure. Women, particularly premenopausal women, are often excluded from clinical studies for safety reasons and due to potential confounds arising from normal cyclic hormonal fluctuations or hormonal contraceptive use [64; 65]. Although there are well-known differences in both metabolic physiology [66–68] and the response to disease [31]; there remains a significant gap in the development of translationally relevant obesogenic models in female animals. Historically, male animals are almost exclusively used in preclinical research; when both sexes are included, they are often underpowered to detect an effect of sex or fail to account for the underlying differences between the sexes in experimental design. In obesity and metabolic disease research, this has led to the prevailing belief that female mice are resistant to diet-induced obesity and thus a poor experimental model. Here, we address this by refining a commonly used paradigm for DIO – 8-12 weeks of HFD feeding – to better optimize it for reliably inducing obesity in female mice. We also investigated the impact of age, sex, and diet on the function of ARH AgRP/NPY neurons and the hypothalamic glial response.

In lean mice, the baseline activity of AgRP/NPY neurons is significantly influenced by sex. Nearly all studies on the physiology of these neurons comes from experiments using exclusively male mice. In *ex vivo* and *in vivo* experiments, in male mice, AgRP/NPY neurons have a robust, highly stereotyped response to metabolic state, with low levels of activity in satiated mice and elevated activity following food deprivation. We show here that this pattern is different in AgRP/NPY neurons from young (4 months old) female mice. In these animals, the activity of these neurons is significantly higher even at baseline in brain slices from lean, satiated female mice compared to those from age-matched male mice. Thus, although there is a slight increase in their output following food deprivation, relative to the baseline level of activity, this is significantly lower than that observed in male mice, suggesting that the fundamental physiological function of AgRP/NPY neurons is strongly sexually dimorphic. As in male mice, we observed an HFD-associated increase in AgRP/NPY neuronal activity, but again, relative to the baseline level, this was less pronounced than previous observations in male mice [3; 24]. We previously reported that chronic (8 weeks) of HFD feeding was associated with both obesity and the development of AgRP/NPY neuronal leptin resistance [3; 69]. Again, this response appears to be strongly sex dependent, as neurons from female mice remain leptin-sensitive after 8 weeks of HFD and an altered response to leptin is not the driving factor behind the sex- and diet-associated changes in AgRP/NPY neuronal function.

Age significantly impacts the excitability of AgRP/NPY neurons in female mice. In middle-aged females, the baseline activity of AgRP/NPY neurons was significantly lower than in younger female mice and is almost identical to that measured by us and others in male mice [3; 24; 25; 70]. As a result, the impact of both fasting and HFD relative to baseline is much more pronounced, perhaps not coincidentally, at the age when female mice begin to develop diet-induced obesity. In humans, the drop in ovarian estrogen during perimenopause causes changes in lipid metabolism, energy balance, insulin resistance and body fat distribution, with postmenopausal women developing more ‘male-like’ patterns compared to pre-menopausal women [71]. The transition to menopause may be influenced by obesity, with reports that women with obesity have a later age of onset, possibly because of pro-estrogenic effects of the expanded adipose tissue [71; 72]. Female mice also undergo ovarian aging and experience a transition known as ‘estropause’, although unlike humans, mice do not experience the large drop in estrogen production, though the changes in estrous cycle regularity, ovarian reserve, cellular senescence, inflammation, and fertility are similar to those observed in peri- and post-menopausal women. In mice, estropause generally occurs around 9-12 months [73; 74], within the window used here as our ‘middle-aged’ group, so it is tempting to speculate that the emergence of diet-induced weight gain and associated changes in hypothalamic neuronal function and inflammatory markers represents an interaction between the decreased ovarian function in aging female mice and consumption of a calorie-dense high-fat diet.

In mammals, there is tight regulation between reproduction and metabolism to ensure that females are able to maintain pregnancy and support offspring [75–77]. Consistent with their critical role in regulating energy homeostasis, AgRP/NPY neurons play a central role in linking metabolic function to reproductive fertility [78–81]. Thus, it is not surprising that there is a pronounced sex dimorphism in the function of these neurons and their response to an environmental dietary factor, particularly in young, fertile mice. Although all of the mice used here were virgin animals, they were gonadally intact and their regulation of the hypothalamic-pituitary-gonadal axis was functioning normally, so our observations here may reflect a normal element of the role of AgRP/NPY neurons in both female reproduction and its relationship to metabolic health.

In summary, we show that the known sexual dimorphism in the development of obesity may arise in part from sex-dependent differences in the function and regulation of AgRP/NPY neurons in the ARH. These neurons play a central role in the central melanocortin system, which is essential for the regulation of appetite, satiety, and body weight, as well as the integration of both central and peripheral signals for homeostatic maintenance. There are well-described sex differences in metabolism, so it is perhaps not surprising that there are also significant differences in the function of these neurons and in the way they respond to a HFD and the development of obesity. Our results highlight the critical need to include both sexes to understand both fundamental aspects of physiology and how sex differences in these pathways influence susceptibility for disease, including obesity and metabolic syndrome.

## Acknowledgements

This work was funded by NIH R01DK102918, RF1AG059778 to K.M.S.O. and NIH F32DK120298 to A.C.K..

## CRediT authorship contribution statement

**Tionna Brothers -** formal analysis, investigation, visualization, writing – original draft; **Wei Wei** – formal analysis, investigation, writing – review and editing; **Austin C. Korgan –** formal analysis, investigation, methodology, writing – review and editing; **Zoey Bridges –** investigation, writing – review and editing; **Kristen M.S. O’Connell –** Conceptualization, data curation, formal analysis, funding acquisition, methodology, supervision, visualization, writing – original draft.

